# Presynaptic HCN channels constrain GABAergic synaptic transmission in pyramidal cells of the medial prefrontal cortex

**DOI:** 10.1101/2021.05.23.445332

**Authors:** Wei Cai, Shu-Su Liu, Bao-Ming Li, Xue-Han Zhang

## Abstract

Hyperpolarization-activated cyclic nucleotide-gated (HCN) channels are widely expressed in neurons in the central nervous system. It has been documented that HCN channels regulate the intrinsic excitability of pyramidal cells in the medial prefrontal cortex (mPFC) of rats. Here, we report that HCN channels limited GABAergic transmission onto pyramidal cells in the mPFC. Pharmacological block of HCN channels resulted in a significant increase in the frequency of both spontaneous and miniature inhibitory postsynaptic currents (IPSCs) in mPFC pyramidal cells. Such facilitation effect on mIPSCs required presynaptic Ca^2+^ influx and reversed by high-dose cAMP. Such facilitation did not exist in the presence of the T-type Ca^2+^ channel selective blockers. Immunofluorescence staining revealed that HCN channels expressed in presynaptic GABAergic terminals, as well as in both soma and neurite of parvalbumin-expressing (PV-expressing) basket cells in the mPFC. The present results indicate that HCN channels in GABAergic interneurons, most likely PV-expressing basket cells, constrain inhibitory control over layer 5-6 pyramidal cells through restricting presynaptic Ca^2+^ entry.

## Introduction

Hyperpolarization activated cyclic nucleotide-gated (HCN) channels are richly expressed in the central nervous system, which are consist of four either identical or nonidentical subunits (HCN1-4)[1], are activated with membrane hyperpolarization, and are regulated directly by cAMP [2–4]. HCN channels conduct a current called I_h_, which contributes to resting potential and input resistance. HCN channels have a major role in controlling neuronal intrinsic excitability, dendritic integration of synaptic potentials, synaptic transmission, and rhythmic oscillatory activity in individual neurons and neuronal networks [4–9].

HCN channels are principally located in pyramidal cell dendrites, although they are found at lower densities in the soma of pyramidal neurons as well as other neuron subtypes[10]. Somato-dendritic HCN channels in pyramidal neurons modulate spike firing and synaptic potential integration by influencing the membrane resistance and resting membrane potential[10]. In addition to their dendritic localization, HCN channels are expressed in cortical and hippocampal axons and synaptic terminals of inhibitory and excitatory neurons [5, 9, 11–13]. Additionally, presynaptic HCN current, I_h_, has been indicated to affect excitatory synaptic transmission in invertebrate neurons and vertebrate neurons where Ih has been shown to influence excitatory transmitter release [14–17] via affecting the activities of presynaptic terminal Ca ^2+^ channels [17]. Additionally, presynaptic Ih affect inhibitory neurotransmission in the rodent globus pallidus, cerebellum and hippocampus [12, 13, 18, 19], though the mechanism by which this occur is unknown.

The prefrontal cortex (PFC) plays a critical role in cognitive functions. The PFC is composed of two major neuronal populations: glutamatergic pyramidal neurons and γ-aminobutyric acid (GABAergic) interneurons. Although GABAergic interneurons only account for approximately 20% of the cortical neuronal population, they are critical elements of cortical circuits by providing feedforward and feedback inhibition and generating synchronous and rhythmic network activity [20].

Although GABAergic interneurons are a minority of the neuron population in the PFC (10%-20%), each interneuron could control hundreds to thousands of pyramidal cells through its profuse local axonal arborizations. Somatostatin, Calretinin, Parvalbumin (PV)-expressing basket cells comprise ~50% of GABAergic neuron population in the neocortex [21]. Axons of PV-expressing basket cells preferentially target the soma and proximal dendrites of pyramidal cells, forming multiple inhibitory synapses with a high probability of GABA release [21–24]. Thus, PV-expressing basket cells exert a powerful inhibitory control over pyramidal cells and are likely to constitute the dominant inhibitory system in the PFC.

It has been reported that peri somatic inhibition ensured by basket cells also has a powerful regulatory effect on the synchronization and oscillation of pyramidal cells [22, 25]. Neuronal synchronization and oscillation are necessary for the execution of cognitive functions [26], and abnormal synchronization and oscillation in the PFC may result in cognitive deficits seen in psychiatric disorders [27]. Wang et al. reported that the α_2A_ adrenoceptor-cAMP-HCN channel signaling pathway in monkey prefrontal cortical cells plays an important role in maintaining the delay-period persistent firing, as such facilitating working memory, although the cell-type localization of HCN channels remains to be identified [28].

However, little is known about the role of HCN channels in GABAergic interneurons of the cortex in GABAergic transmission. The present study attempted to examine if and how HCN channels in interneurons regulates inhibitory synaptic transmission onto layer 5-6 pyramidal cells in the medial prefrontal cortex of rats, using immunofluorescence staining and whole-cell recording approaches.

## Results

### HCN channels limit GABAergic transmission onto pyramidal cells

To examine whether HCN channels are involved in regulating GABAergic synaptic transmission in the mPFC, we recorded action potential-dependent spontaneous IPSCs (sIPSC) in layer 5-6 pyramidal cells in the presence of 20 μM DNQX and 50 μM D-APV with −70 mV holding potential (Fig. 1A). The recorded sIPSCs could be completely blocked by the GABA_A_ receptor antagonist bicuculline (10 μM) (data not shown). Under the control experimental condition, the frequency of sIPSC, especially, large sIPSCs (amplitude >20pA) was 1.95±0.37 Hz, and it increased to 3.28±0.5 Hz 12-15 min after bath application of HCN channel blocker ZD7288 (30 μM) (Fig. 1C; *p*<0.01, *paired t*-test, n=6 cells). The facilitation effect of ZD7288 was largely reversible 12-15 min after termination of ZD7288 (Fig. 1C; 2.62±0.31 Hz after washout). The enhancement of the frequency of large sIPSCs in pyramidal cells indicates that HCN channels limit GABAergic transmission onto pyramidal cells.

**Figure 1.**
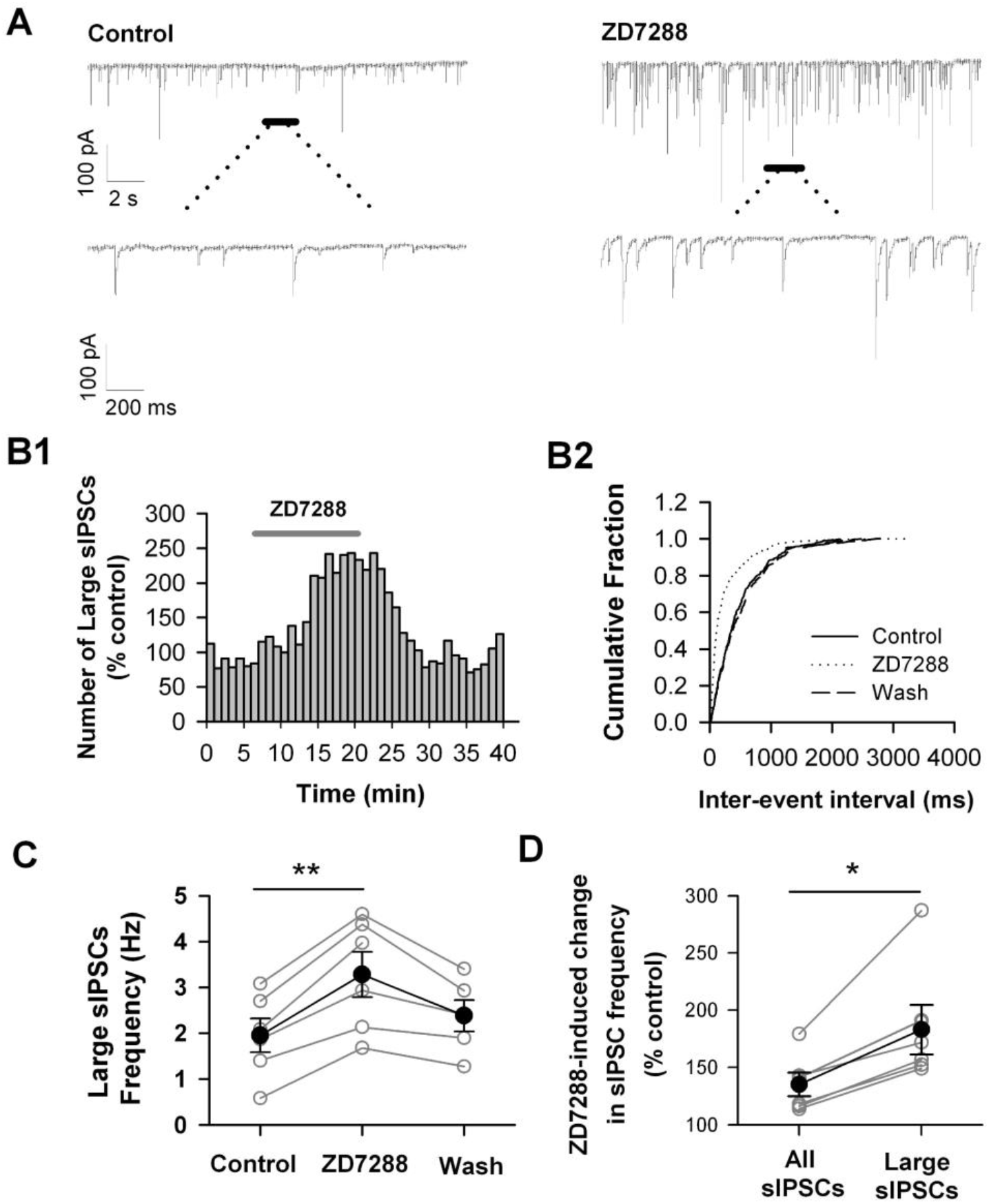
Blockade of HCN channels increases the frequency of sIPSCs. *A*. An example trace of spontaneous IPSCs (sIPSCs) recorded in mPFC pyramidal cell in absence (*Control*) or presence of HCN channel blocker ZD7288 (30 μM). Holding potential = − 70 mV. B. ZD7288 increases the frequency of sIPSCs with large amplitude (> 20 pA). The number distribution of large sIPSCs (bin=60 s; *B1*), and the cumulative fraction distribution of inter-event intervals of sIPSCs before (*Control*), during (*ZD7288*), and after ZD7288 application (*Wash*; *B2*). Data were from the same cell in (A). C. The summary individual (open circles) and grouped (closed circles) frequency of large sIPSCs. n=6 cells. \*\**p*< 0.01 D. The sIPSC frequency changes in all detective events and large events after ZD7288 application. Open circles for the individual cell; Close circles for grouped cells. Data were from the same cell in (A). \**p*<0.05

### Presynaptic but not postsynaptic HCN channels are involved in limiting GABAergic transmission

To test HCN channels affect GABAergic transmission by postsynaptic or presynaptic pathway, we test miniature IPSCs(mIPSC), which can represent responses of pyramidal cells to spontaneous release of single GABA-containing vesicles and action potential independently. Therefor mIPSCs were recorded in the presence of 1 μM tetrodotoxin (TTX) that blocks action potential firing and propagation. As shown in Figure 2, the frequency of mIPSCs was 1.37±0.20 Hz and the peak amplitude was 13.21±1.73 pA under control condition. When ZD7288 (30 μM) was applied, the frequency of mIPSCs increased to 2.40±0.39 Hz (Fig. 2D1; *p*<0.01, *paired t*-test, n=10 cells), whereas the amplitude of mIPSCs kept unchanged (Fig. 2D2; control: 13.22±1.59 pA, ZD7288: *p*>0.05, *paired t*-test). The facilitation effect of ZD7288 on mIPSC frequency was largely recovered after termination of ZD7288 application (2.08±0.30 Hz after washout; Fig. 2D1). The fast rise time of mIPSCs in the presence of ZD7288 was comparable with control (10%-90% rise time: 1.52±0.12 ms under control and 1.54±0.13 ms in the presence of ZD7288; *p*>0.05, *paired t*-test; Fig. 2E). Thus, ZD7288 did not change the kinetics of mIPSCs. Together, the mIPSC events regulated by ZD7288 were mainly generated in the presynaptic region of recorded pyramidal neurons [29]. These results suggested that HCN channels may limit presynaptic GABA release to constrain GABAergic transmission onto pyramidal cells.

**Figure 2.**
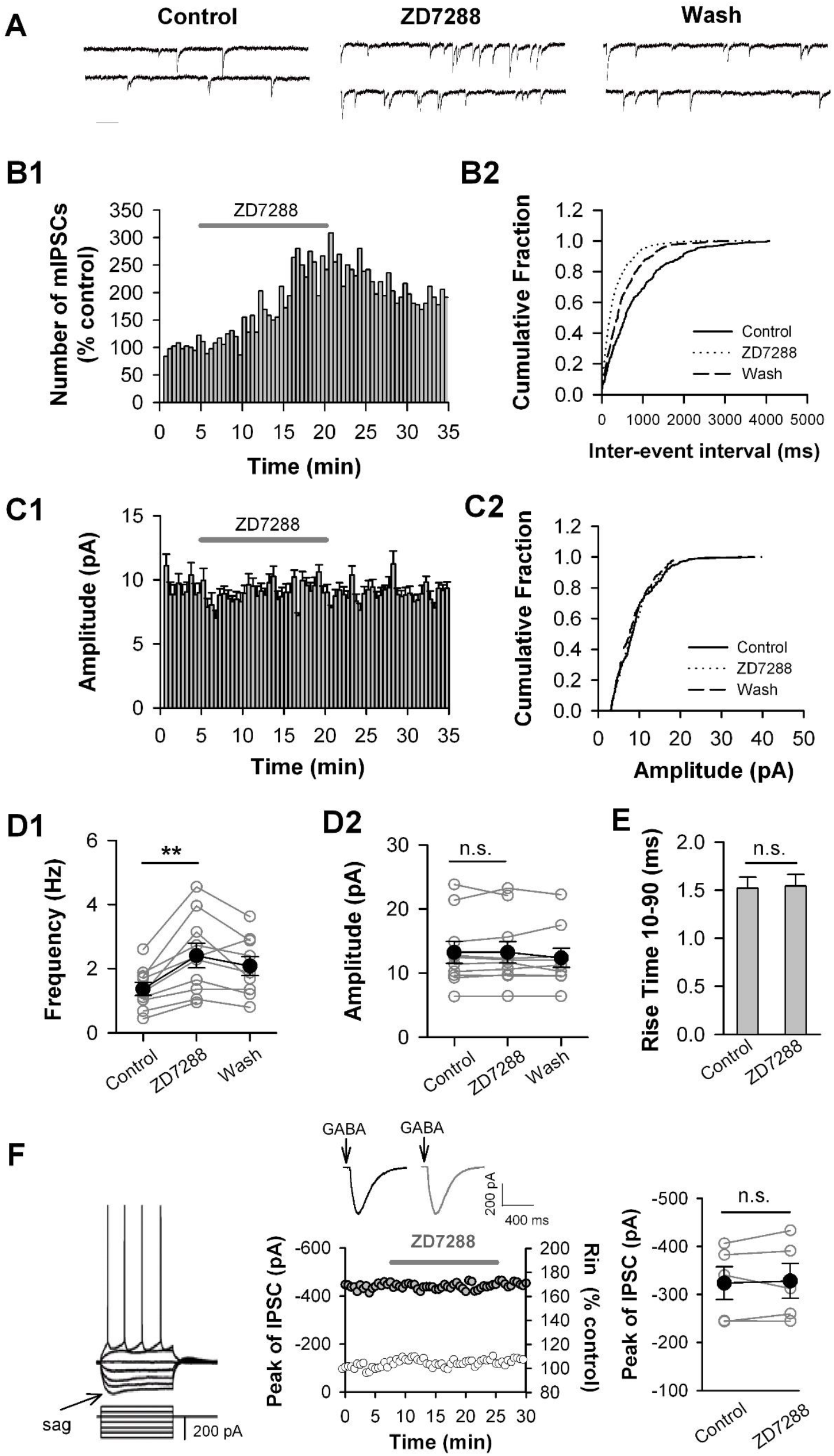
Blockade of HCN channels enhances the frequency but not amplitude of mIPSCs. *A.* Representative traces of miniature IPSCs (mIPSCs) recorded in mPFC pyramidal cell before (*Control*), during (*ZD7288*), and after ZD7288 application (*Wash*). Holding potential: −70 mV. Calibration: 20 pA, 200 ms. B. ZD7288 facilitates the frequency of mIPSCs. The number distribution of mIPSCs (bin=60 s, *B1*), and the cumulative fraction distribution of inter-event intervals of mIPSCs (*B2*). Data were from the same cell in A. C. ZD7288 has no effect on the amplitude of mIPSCs. The amplitude distribution of mIPSCs (bin=60 s, C1), and the cumulative fraction distribution of mIPSC amplitude (C2). Data were from the same cell in A. D. The mIPSC frequency (D1) and amplitude (D2) from the individual cell (open circles) and grouped cells (closed circles). n=10 cells, \*\**p*<0.01. E. ZD7288 has no effect on 10-90% rise time of mIPSCs. Data were from the same cells in D. F. ZD7288 has no effect on the amplitude of IPSCs evoked by puff application of GABA (10 μM) to pyramidal cells. A typical example for pyramidal cells with a sag (*left*). An example time course of the IPSC amplitude (black circles) and the input resistance (grey circles) obtained from the pyramidal cell. The insets show the IPSC traces in the absence (black) and presence of ZD7288 (grey), each of which is the average of seven consecutive IPSCs (*middle*). The summary individual (open circles) and grouped (closed circles) amplitude of IPSCs (*right*). n=5 pyramidal cells.

To clarify whether HCN channels in postsynaptic pyramidal cells are involved in the ZD7288 facilitation effect on GABAergic transmission, we evoked postsynaptic GABA-receptor-mediated currents by puffing GABA onto the soma of pyramidal cells, and examined the effect of ZD7288 on GABA-evoked currents. In this experiment, we only examined the effect of ZD7288 on GABA-evoked currents in the pyramidal cells with depolarizing sag to ensure the recorded pyramidal cells expressing HCN channels (Fig. 2F, left). It is well known that the depolarizing sag in response to negative current injection is a typical of response pattern in pyramidal cells expressing HCN channels in the PFC [8] and other tissues [30–32]. GABA-evoked currents recorded in the presence of TTX (1 μM), DNQX (20 μM) and D-APV (50 μM) with −70 mV holding potential (Fig. 2F, *middle, inset*). Puff application of GABA (10 μM) evoked an outward current that was completely blocked by the GABA_A_ receptor antagonist bicuculline (10 μM; n=3; data not shown). ZD7288 had no effect on the amplitude of GABA-evoked currents (Fig.2F, *right*; Control: −323.59±34.04 pA; ZD7288: −328.10±36.71 pA; ZD7288 did not change the input resistance (Fig. 2F, *middle*; 108.55±3.10% of control 12-15 min after ZD7288 application, p>0.05 for ZD7288 vs. control, paired t-test). Thus, the ZD7288-induced increase in the frequency of mIPSC was not due to the blockade of HCN channels in the pyramidal cells, but most likely resulted from the blockade of HCN channels in GABAergic terminals innervating the pyramidal cells.

### HCN channels are present on presynaptic GABAergic terminals

To identify that the expression of HCN channels in presynaptic GABAergic terminals, we examined the expression of HCN channels in GABAergic terminals using immunostaining technique. We performed immunolabeling against GAD65, the synthetic enzyme for GABA, to label GABAergic terminals[33]. We double labeled HCN1, HCN2, and HCN4 channels with GAD65. Single-plane confocal image showed that GAD immunoreactive (GAD-ir) appeared in puncta structures distributed in the neuropil, as well as around unlabeled cell bodies (Fig. 4C), which consistent with previous reports [34]. Merging single-plane images showing HCN1-ir, HCN2-ir, and HCN3-ir with showing GAD65-ir showed the puncta of GAD65-ir, surrounding the cell bodies of HCN-ir cells, partially co-located with HCN1-ir, HCN2-ir, and HCN3-ir (Fig. 4D). These data point out that HCN channels are present in the GABAergic terminals, indicating that blockade of HCN channels affects presynaptic GABA release.

### HCN channel activation suppress GABA release

Some researchers have pointed out that ZD7288 can activate Na^+^ and Ca^2+^ channels[35, 36]. Thus, we ought to clarify this phenomenon is based on HCN channels. Activation of HCN channels is facilitated by cAMP. cAMP strongly enhances HCN channel function [37–41]. In cortex, HCN channels are heteromers of HCN1 and HCN2 subunits that are highly responsive to cAMP [38, 39]. We hypothesized that up-regulation of presynaptic HCN channel function might alter GABA release onto pyramidal cells. To test this hypothesis, the frequency of mIPSC was compared after perfusion of the membrane-permeable, cAMP analog, Sp-cAMPs (200 μM). Perfusion of Sp-cAMPs (5 min) markedly decreased the frequency of mIPSCs in pyramidal cells. Grouped data demonstrate that Sp-cAMPs (12-15 min after application) significantly decreased mIPSC frequency from 1.71 ±0.28 Hz to 1.26 ±0.24 Hz (p<0.01, paired t-test; n=5 pyramidal cells from 2 animals; Fig. 4C). The effect of Sp-cAMPs on mIPSC frequency was largely recovered after termination of Sp-cAMPs application (1.56 ±0.26 Hz after washout; p<0.05 vs. Sp-cAMPs treatment; paired *t*-test; Fig. 4C). This result solid our hypothesis that ZD7288 influence mIPSC through HCN channels.

### HCN-channel blockade facilitates GABA release via presynaptic Ca^2+^ influx

It has been proved that Ca^2+^ influx is critical for the vesicle releasing at presynaptic membrane[42]. To examine whether Ca^2+^ influx is involved in the ZD7288-induced facilitation of mIPSC frequency, we first examined the effect of ZD7288 under Ca^2+^-free condition. As shown in Figure 3, ZD7288 failed to increase mIPSC frequency in the absence of extracellular Ca^2+^. The mIPSC frequency 12-15 min after ZD7288 application was 106.4 ±5.27 % of control (*p*>0.05, *paired t*-test, n=5 pyramidal cells from 4 animals; Fig. 5A). We next investigated the effect of ZD7288 in the presence of Cd^2+^ (200 μM), a Ca^2+^ channel blocker. ZD7288 significantly increased mIPSC frequency (163.48 ±10.05 % of control, *p*<0.01 for ZD7298 *vs*. control, paired t-test; Fig. 5B). Such facilitation was completely blocked when Cd^2+^ was added into perfusion solution (108.0 ±5.82 % of control; *p*<0.01 for ZD+Cd^2+^*vs*. ZD alone, paired t-test, n=6 pyramidal cells from 3 animals). Thus, these data indicate that Ca^2+^ influx is required for the ZD7288-induced augmentation of mIPSC frequency.

**Figure 3.**
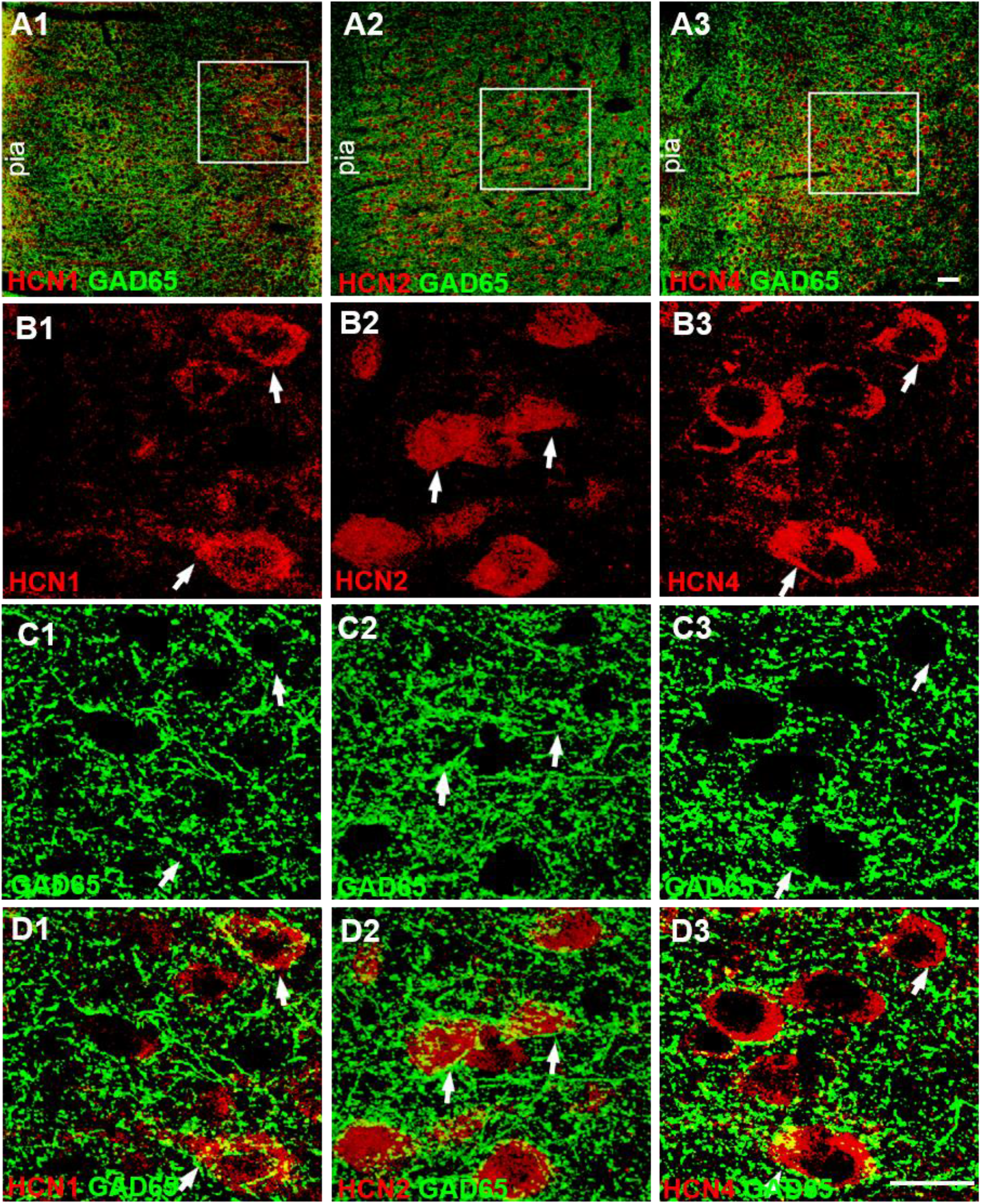
HCN channels are present on GABAergic terminals in the mPFC. **A.** Low-magnification confocal images showing double stained with HCN channels (red) and GAD65 (green), a GABAergic terminal marker. The squares illustrating the cells in layer 5-6 of the mPFC. Scale bar, 40 μm **B** and **C.** Single-plane confocal images showing the HCN1-ir (*B1*), HCN2-ir (*B2*), HCN4-ir (*B3*), and GAD65-ir (*C1-C3*) at high magnification. GAD65-ir appears in punctuate structures distributed in the neuropil, as well as around unlabeled pyramidal cell soma (C1-C3). E. Merging of the paired images (B1 and C1), (B2 and C2), and (B3 and C3) shows that the puncta of GAD65-ir surround the cell bodies of HCN1-ir (B1), HCN2-ir (B2), and HCN4-ir (B3) cells. Partially overlapping areas of red (HCN) and green (GAD65) profiles showing yellow. The arrowheads indicate double-labeled cells. Scale bars represent 20 μm.

**Figure 4.**
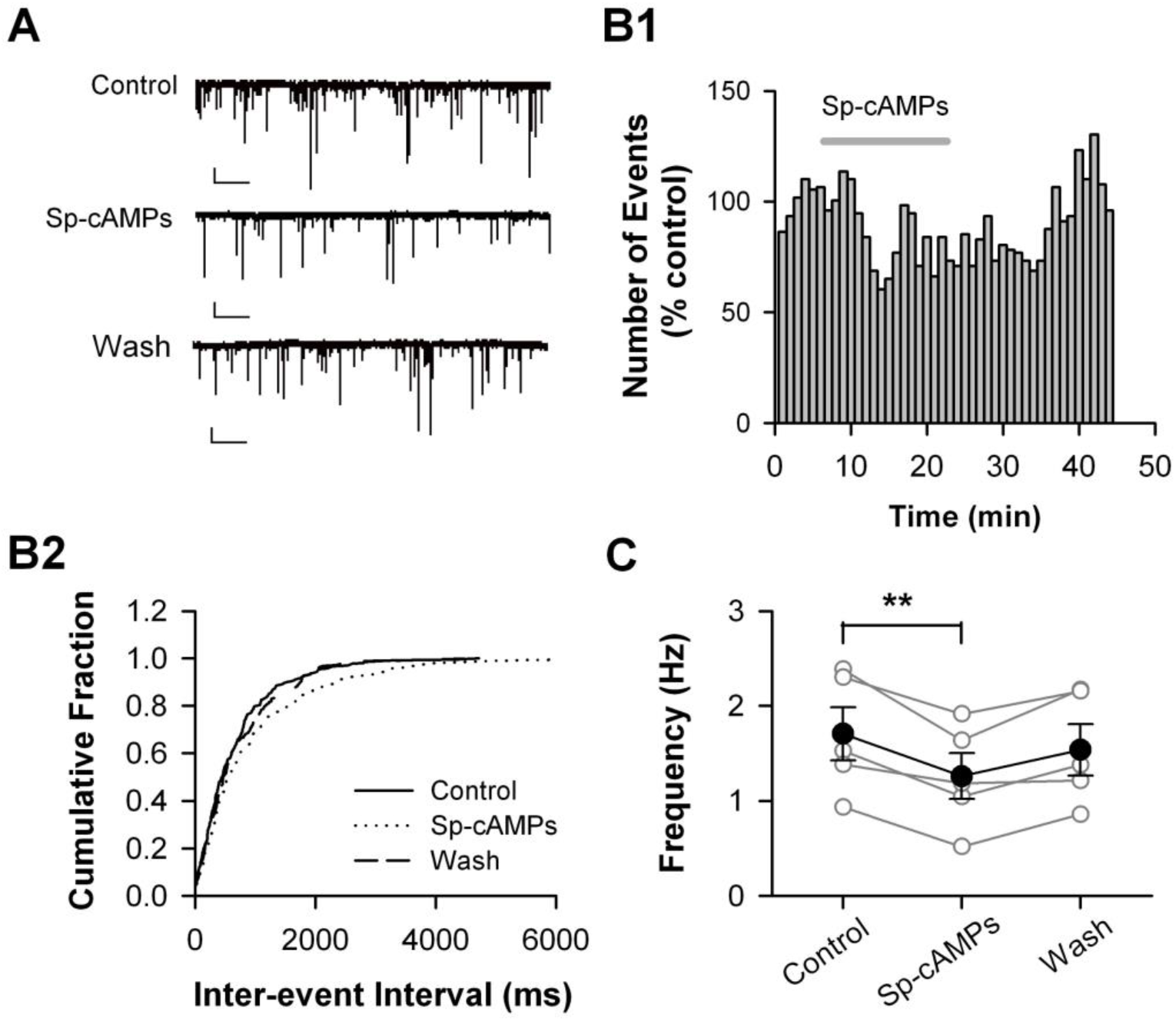
Enhancing HCN channel function decreases the frequency of mIPSC. *A.* Representative traces of miniature IPSCs (mIPSCs) recorded in mPFC pyramidal cell before (*Control*), during (*Sp-cAMPs*), and after Sp-cAMPs application (*Wash*). Holding potential: −70 mV. Calibration: 20 pA, 200 ms. B. Sp-cAMPs decreases the frequency of mIPSCs. The number distribution of mIPSCs (bin=60 s; *B1*), and the cumulative fraction distribution of inter-event intervals of mIPSCs (*B2*). Data were from the same cell in A. C. Summary for individual cell (open circles) and grouped cells (closed circles). n=5 cells, \*\**p*<0.01.

**Figure 5.**
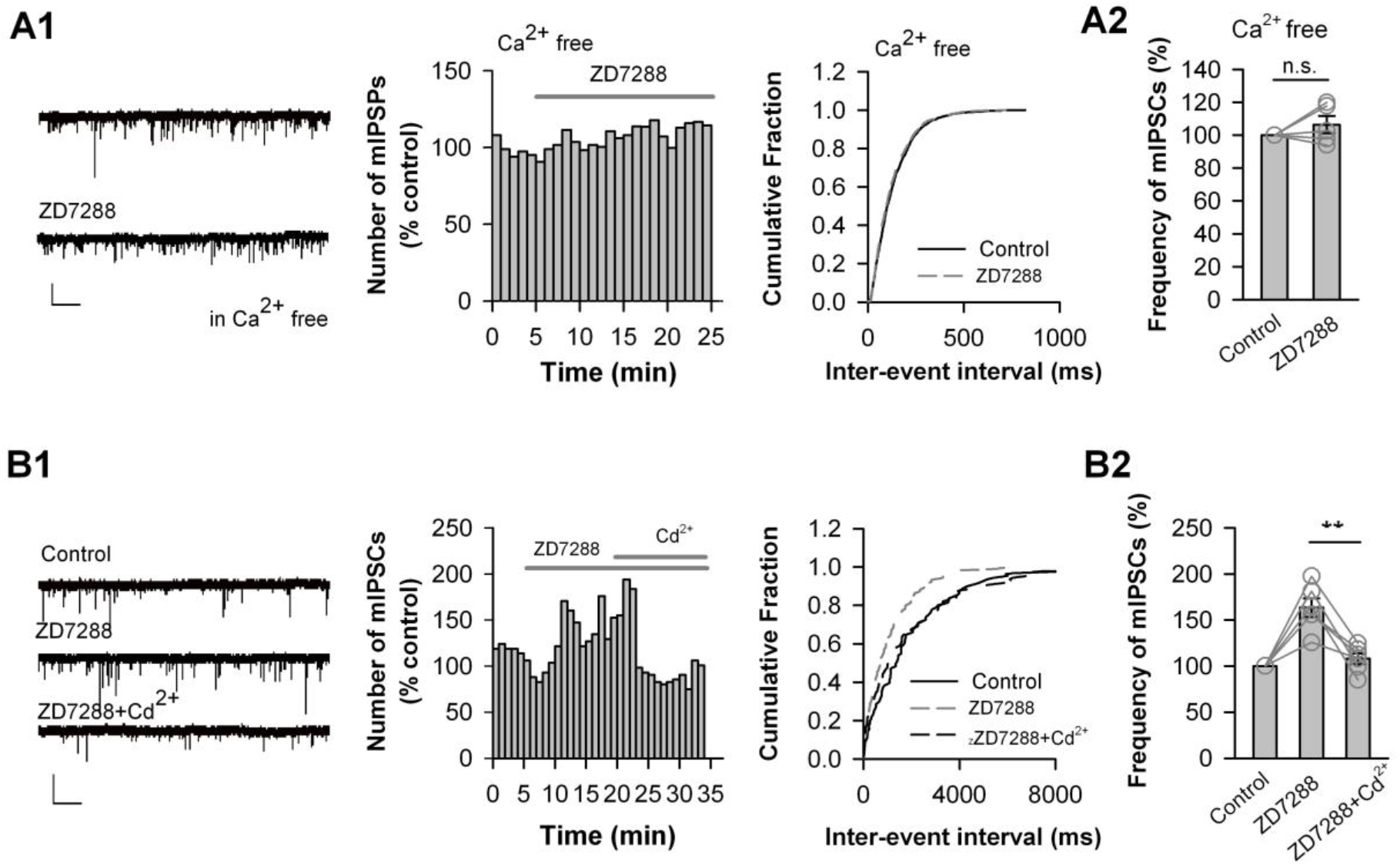
HCN-blockade enhancement of mIPSC frequency requires Ca^2+^ influx. A. ZD7288 has no effect on mIPSC frequency under the condition of omitting extracellular Ca^2+^. An example trace of miniature IPSCs (mIPSCs) recorded in pyramidal cell under Ca^2+^-free perfusion solution (*A1, left*). The number distribution of mIPSCs (bin=60 s; *A1, middle*), and the cumulative fraction distribution of inter-event intervals of mIPSCs (*A1, right*) recorded from cell in left. The individual and grouped data showing the change in mIPSC frequency induced by ZD7288 under extracellular Ca^2+^-free condition. n=5 pyramidal cells. B. Blocking Ca^2+^ channel abolishes the effect of ZD7288 on mIPSC frequency. An example trace of miniature IPSCs (mIPSCs) recorded in pyramidal cell (*B1, left*). The number distribution of mIPSCs (bin=60 s; *B1, middle*), and the cumulative fraction distribution of inter-event intervals of mIPSCs before (*control*), during application of ZD7288 alone (*ZD7288*), and during co-application of ZD7288 and Ca^2+^ channel blocker Cd^2+^ (200 μM, *ZD7288*+*Cd*^2+^) (*B1, right*) recorded from cell in *left.* The individual and grouped data showing the changes in mIPSC frequency induced by ZD7288 alone, and co-application of ZD7288 and Cd ^2+^ (*B2*). **p<0.01, n=6 pyramidal cells. Calibrations: 5s, 20 pA in both A and B.

### Voltage-gated Ca^2+^ channels are involved in facilitation of GABA release by HCN blocking

How loss the function in HCN channels increases Ca^2+^ influx? Considering that blockade of HCN channels hyperpolarizes membrane potential [3, 8], a possibility is that hyperpolarization of resting membrane potential results in voltage-sensitive Ca^2+^ channel (VGCC) open and thereby increasing Ca^2+^ influx into presynaptic terminals. To address this possibility, we hyperpolarized resting membrane potential by reducing external K^+^ concentration from 2.5 to 1.75 mM, which, according to the Nernst equation, could hyperpolarize the resting membrane potential by ~10 mV. As shown in Fig. 6A, such a reduction in external K^+^ concentration resulted in hyperpolarization of membrane potential by 7.0 ±3.0 mV (n=6 pyramidal cells), and induced an upregulation in mIPSC frequency by 168.94 ±7.28% of control (*p*<0.01, paired t-test; n=5 pyramidal cells from 3 animals). Thus, hyperpolarizing membrane potential mimics the effect of blocking HCN channels on mIPSC frequency.

**Figure 6.**
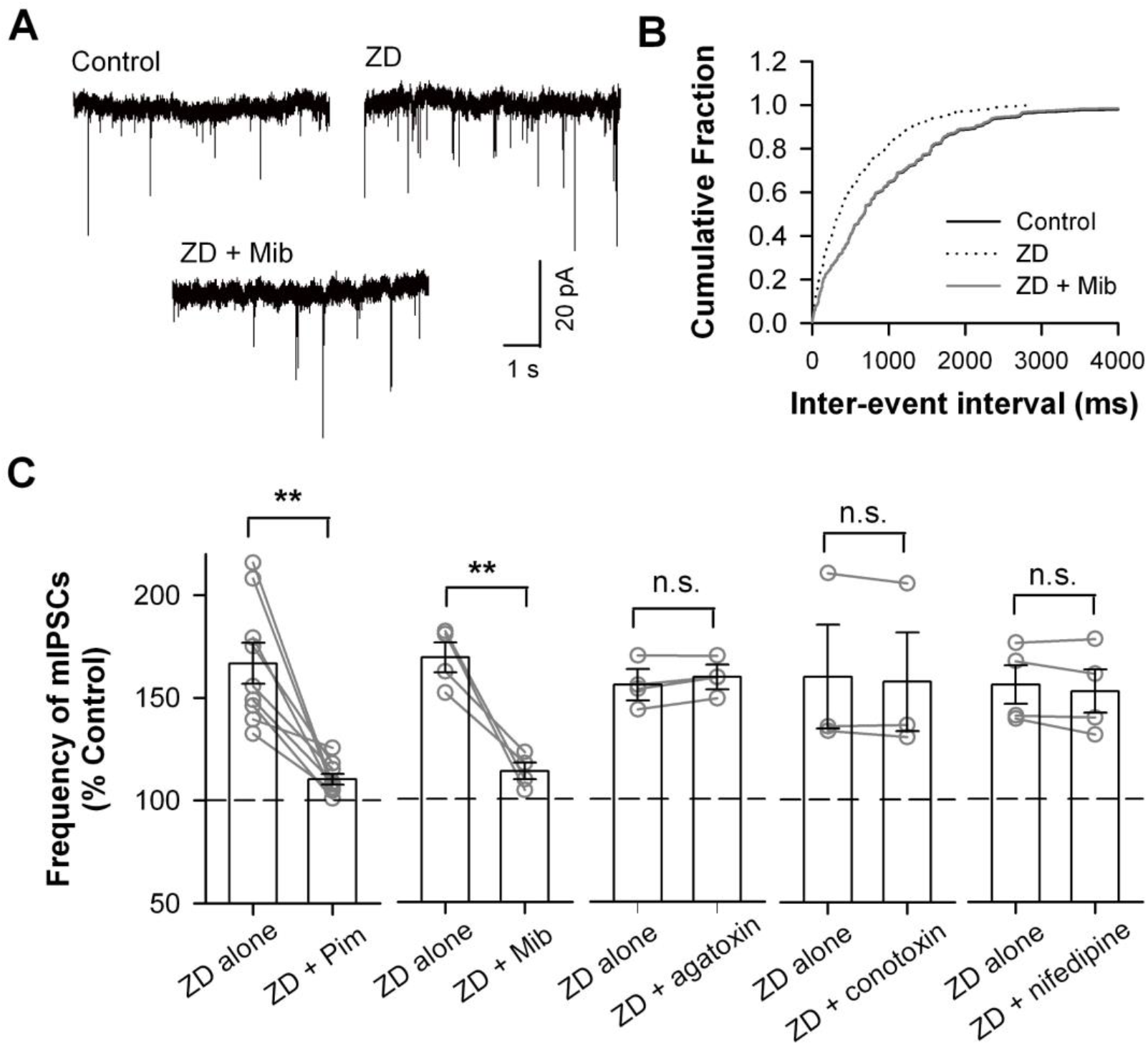
T-type Ca2+ channel blockers occlude the increment in mIPSC frequency induced by blocking HCN channels. **A.** Representative traces of mIPSCs recorded in pyramidal cell (*left*), and the cumulative fraction distribution of inter-event intervals of mIPSCs before (*Control*) and 8-10 min after hyperpolarization. Hyperpolarization of resting membrane potential by reducing external K^+^concentration from 2.5 mM (control) to 1.75 mM. Holding potential: −70 mV. Calibration: 1 s, 20 pA. **p<0.01. n=4 pyramidal cells. **B.** the cumulative fraction distribution of inter-event intervals of mIPSCs recorded in pyramidal cell before (*Control*), during (*ZD7288*), and after co-application of ZD7288 with T-type Ca^2+^ channel selective blocker mibefradil (Mib; 10 μM) (*ZD+Mib; left*), Holding potential: −70 mV. **C.** Bar graph demonstrating the effects of co-application of ZD7288 (30 μM) and Ca2+ channel blockers for T-type (pimozide, 1 μM; mibefradil, 10 μM), P/Q-type (ω-agatoxin IVA, 500 nM), N-type (ω-Conotoxin GVIA, 500 nM), and L-type (nifedipine, 2 mM) Ca2+ channels. Open circles for individual cells and bar for grouped data. **p<0.01, paired t-test.

Next step we want to figure out which type of VGCCs are involved in Blockade of HCN channels changes resting membrane potential and alters the activity of voltage-gated Ca^2+^channels (VGCC), such as low voltage-activated T-type or high voltage-activated P/Q-type, N-type, and L-type Ca^2+^ channels, resulting in Ca^2+^ influx into presynaptic terminals innervating pyramidal cells, and an increase in GABA release from the terminals.

HCN channels open at membrane potentials more negative to −50 mV and are important for regulating resting membrane potential. HCN channels are also permeable to Na^+^ and K^+^ and form an inward current at rest, thereby depolarizing resting membrane potential. Thus, it is possible that blockade of HCN channels changes resting membrane potential and alters the activity of low-voltage-activated T-type or high-voltage-activated P/Q-type, N-type, and L-type Ca^2+^ channels, resulting in Ca^2+^ influx into presynaptic terminals innervating pyramidal cells, and an increase in GABA release from the terminals. To test this possibility, we examined whether the ZD7288-effect on mIPSC frequency could be occluded by Ca^2+^ channel blockers. As shown in Fig. 6, ZD7288 had on effect on mIPSC frequency in the presence of T-type Ca^2+^ channel blocker pimozide (1 μM)[43] (Fig. 6B; *p*<0.01 for pimozide+ZD vs. pimozide, *unpaired t*-test). Additionally, Adding T-type Ca2+ channel blocker mibefradil (Mib; 10 μM)[44] into perfusion solution occluded the facilitation effect of ZD7288 on mIPSC frequency (p>0.05), while perfusing Mib (10 μM) alone did not affect the frequency of mIPSC (p>0.05). But the facilitation of ZD7288 on mIPSC frequency still exist in the presence of the P/Q-type Ca^2+^ channel blocker ω-agatoxin IVA (500 nM), the N-type Ca^2+^ channel blocker ω-conotoxin GVIA (500 nM), and the L-type Ca^2+^ channel blocker nifedipine (Fig. 6C; *p*>0.0, *one-way ANOVA*). Together, these results indicate that blockade of HCN channels enhanced GABA release onto pyramidal cells by increasing Ca^2+^ influx through T-type Ca^2+^ channels.

### HCN channels express in parvalbumin-expressing basket cells

To examine the expression of HCN channels in GABAergic interneurons in layer 5-6 of the mPFC, we conducted double-labeling immunofluorescence staining. Four HCN channel subunits, HCN1, HCN2, HCN3, and HCN4, have been cloned that contribute to brain HCN channels [45], and all the four HCN subunits are expressed in the mammalian brain, of which HCN3 exhibits the weakest expression [4]. As results presented in Fig. 1 and 2, blockade of HCN channels induced a dramatic increase in GABA release onto pyramidal cells were most likely generated from the soma of pyramidal neurons [29]. Interneurons expressing Ca^2+^-binding protein parvalbumin (PV) mainly innervate the soma of pyramidal cells and constitute the largest population of interneurons (~ 50%) in the prefrontal cortex [21, 23, 29, 46, 47].

We double labeled HCN1, HCN2, and HCN4 subunit with PV. The confocal microscopy images demonstrated that HCN1-ir, HCN2-ir and HCN4-ir (green) abundantly co-localized with PV-ir (red) (Fig. 7A-C). The high-magnification images depicted that both HCN1-ir and HCN2-ir was observed in neurites of PV-ir cells, whereas no HCN4-ir was observed (Fig. 7D and E). Thus, HCN1, HCN2 and HCN4 subunit are all present in the cell bodies of PV-expressing interneurons, while only HCN1 and HCN2 express in the processing of PV-expressing interneurons in the layer 5-6 of the mPFC.

**Figure 7.**
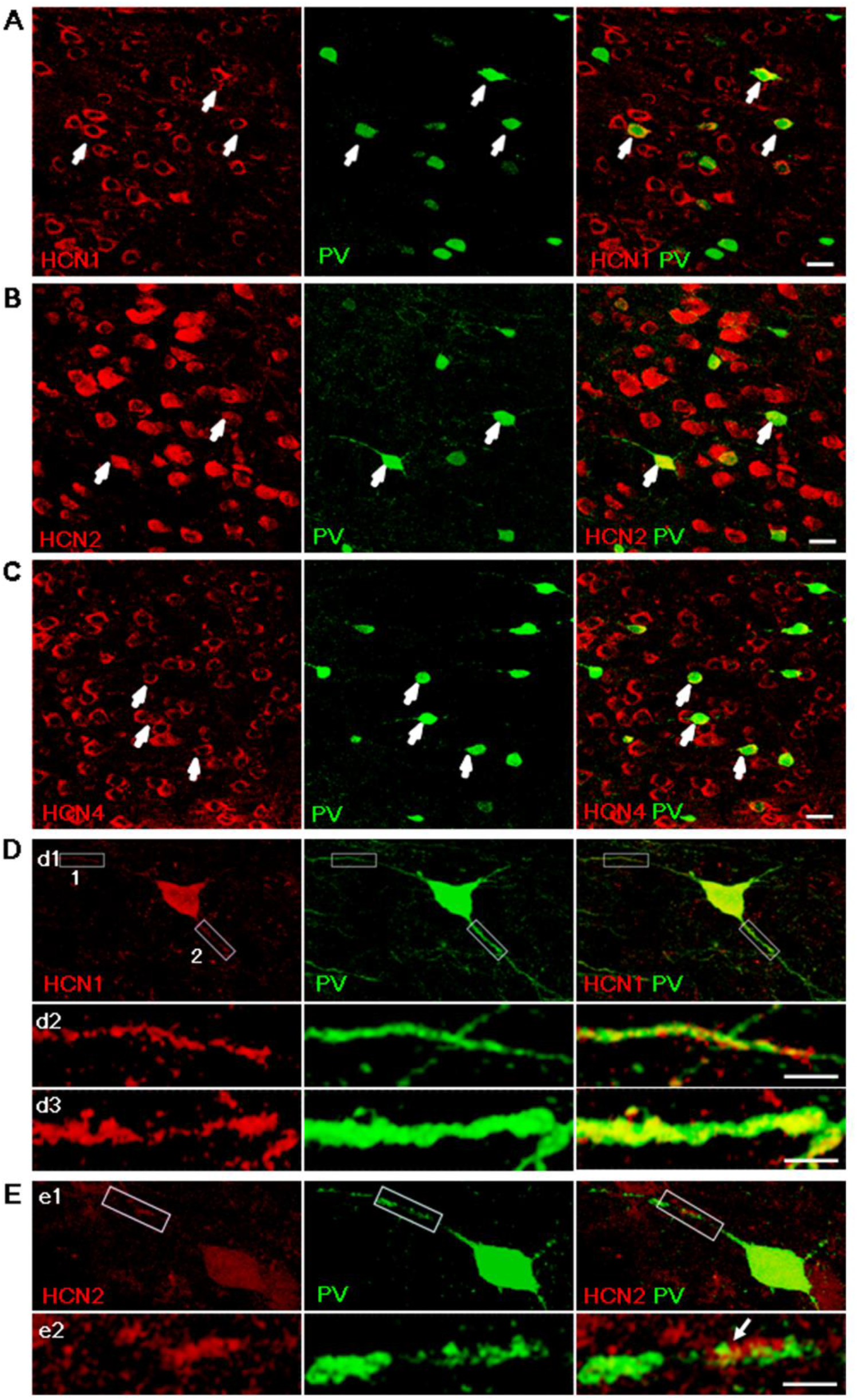
HCN channels are present in soma and neurite of parvalbumin-expressing basket cells in layer 5-6 of mPFC. A-C. Microscopic confocal images showing HCN1-ir (*A*), HCN2-ir (*B*), and HCN4-ir (*C*) locate in PV-ir interneuron in layer 5-6 of mPFC. Double stained with HCN channels (red) and PV (green). Arrowheads indicate double-labeled cells. Scale bar, 20 μm D. High-magnification confocal microscopy images showing that HCN1-ir localize in the soma (*d1*) and along neurite (*d2-d3*) of PV-ir interneuron. Silhouette frame 1 and 2 in neurite (*d1*) is digital magnified for better view of neurite in (*d2*) and (*d3*), respectively. Scale bars, 20 μm in (*d1*) and 1 μm in (*d2*) and (*d3*). E. High-magnification confocal microscopy images showing that HCN2-ir localize in the soma and along neurite of PV-ir interneuron. Silhouette frame in neurite (e1) is digital magnified for better view of neurite in (e2). Scale bars, 20 μm in (e1) and 1 μm in (e2).

## Discussion

In this study, we demonstrated that HCN channels are richly present in cells expressing parvalbumin, and pharmacological blocking HCN channels enhances GABA release onto pyramidal neurons in layer 5-6 of mPFC through increasing Ca^2+^ influx via T-type Ca^2+^ channels.

T-type Ca^2+^ channels are expressed in parvalbumin-expressing basket cells in the 5^th^ and 6^th^ cortical layers [48, 49], suggesting that HCN channels may provide a regulatory mechanism for controlling GABA releases from GABAergic terminals. It is reported that HCN channels regulates T-type Ca^2+^ channel activity in neuronal dendrites in the hippocampus [50] and in layer 3 glutamate terminals in the entorhinal cortex [16, 17]. Consistently, the present study demonstrated that HCN channels are present in the inhibitory presynaptic terminals of layer 5-6 basket cells in the mPFC, and constrain GABA release by restricting Ca^2+^ entry via pre-synaptic T-type Ca^2+^ channels.

The universality and nonselective cation permeability make HCN channels indispensable for cell excitability. Not only in pyramidal neurons, interneurons excitability homeostasis disorder also can trigger psychosis[51, 52]. Simultaneously, HCN channels engage on intracellular cascade reaction, like work as a downstream of some neurotransmitter receptors such as alpha-2 (α_2_) adrenergic receptor, and the location of HCN channels in post-synapse membrane also facilitate they involve in synaptic signaling[28]. Along this principle, the specific expression of HCN channels in parvalbumin-expressing basket cells suggests it may involve in GABA release[53]. Our study consistence with this point and prove this pathway regulate the inhibitory input of layer 5-6 pyramidal neuron in mPFC. Beside alpha-2 (α_2_) adrenergic receptor, HCN channels may also can be regulated by other neurotransmitter receptor such as dopamine receptor through cAMP pathway. Combine these together, some neurotransmitters would regulate GABAergic transmission by means of affecting HCN channels to control cortex rhythmic oscillatory, which is essential for brain functional running.

Regulation of GABA release by HCN channels is diverse, dependent on their locations and proximity to other ion channels. It is reported that, in the entorhinal cortex and globus pallidus, blockade or loss of HCN channels induces an increase in the frequency of sIPSCs and mIPSCs, with no effect on the amplitude of mIPSCs [12, 54], similar with what we found in the present study. However, some other studies showed that pharmacological blockade of HCN channels with ZD7288 results in a reduction, instead of increase, in the frequency of sIPSCs or mIPSCs in the CA1 and DG regions of the hippocampus and in the cerebellum as well [18, 19, 55]. Such discrepancy might be due to that, the presynaptic HCN channels exerts depolarizing influence on GABAergic terminals, and that blockade of HCN channels inhibits neurotransmitters release via hyperpolarization. Indeed, an increase in the external K^+^ concentration from 2.5 to 5.0 mM reverses the inhibitory effect of HCN-channel blockade on mIPSC frequency [18].

Although pyramidal cells in the mPFC receive inhibitory inputs from several types of interneurons [56], the present results suggest that the increase in sIPSC/mIPSC frequency induced by HCN-channel blockade was mainly originated from increased release of GABA from parvalbumin-expressing basket cells. First, as the source of the dominant inhibitory system with the largest population of interneurons in layer 5-6 of the prefrontal cortex [21], parvalbumin-expressing basket cells should inevitably produce largest somatically-recorded IPSCs [23, 47]. Thus, the frequency of sIPSCs with large amplitude should increase dramatically upon GABA release from parvalbumin-expressing basket cells in the presence of ZD7288. Indeed, our data showed that the ZD7288-induced increase in the frequency of large sIPSCs (with amplitude larger than 20 pA) was up to 183.06±21.07 % of control, whereas that of all detective sIPSCs was only 135.34±10.22%. Second, ZD7288 augmented the frequency but not the amplitude or kinetics of mIPSCs (see Figure 2E), suggesting that GABA was released tonically from the GABAergic terminals close to the soma of the pyramidal cells [29]. Third, it has been shown that interneurons that do not express parvalbumin make synapses near the soma of pyramidal cells and utilize N-type Ca^2+^ channels in terminals for GABA release, while parvalbumin-expressing interneurons utilize P/Q-type Ca^2+^ channels in terminals for GABA release [57, 58]. Our data showed that ZD7288 still increased the frequency of mIPSCs in the presence of the N-type Ca^2+^ channel blocker (see Figure 6C). Taken together, we argue that parvalbumin-expressing basket cells produced the major contribution to the ZD7288-induced increase in spontaneous/miniature IPSCs in layer 5-6 pyramidal cells.

Parvalbumin-expressing basket cells in the cortex play an important role in adjusting the gain of synaptic inputs onto and controlling synchronization and excitatory output of pyramidal cells, and through this mechanism, they control both the number of pyramidal cells activated and the firing frequency of the pyramidal cells[59]. Pyramidal cells in layer 5-6 of the mPFC have been suggested to primarily project to subcortical regions to regulate complex motor functions, behavioral arousal and attention [60]. The dynamic modulation of GABAergic inputs to the pyramidal cells by HCN channels in parvalbumin-expressing basket cells may contribute to the regulation of these physiological states. In addition, oscillations occurring in PFC pyramidal cells are essential for behavioral and cognitive functions [61]. Coherent network oscillations, which are facilitated by GABA released onto pyramidal cells, are required for execution of cognitive functions. Abnormal γ-frequency oscillations observed in schizophrenia has been suggested to be due to a reduction in peri-somatic inhibition in PFC pyramidal cells [62]. Thus, HCN channels in parvalbumin-expressing basket cells might be a potential target for drug development for schizophrenia.

## Materials and Methods

### Electrophysiology

Male Sprague-Dawley rats (4-5 weeks old, 80-130 g) were purchased from SLACCAS (Shanghai, China) and were kept in a 12 h light/dark cycle, and food and water were available ad libitum. All experiments were performed in compliance with the Guide for the Care and Use of Laboratory Animals issued by the National Institutes of Health, USA, and were approved and monitored by the Ethical Committee of Animal Experiments at the Fudan University Institute of Neurobiology (Shanghai, China). All efforts were made to minimize the number of animals used and their suffering.

Rats were anesthetized with pentobarbital sodium (40 mg/kg, i.p.) and rapidly decapitated. Brains were quickly removed and immersed in the 0°C artificial cerebrospinal fluid (ACSF) solution containing (in mM): 119 NaCl, 2.5 KCl, 2.5 CaCl2, 1.3 MgCl2, 26.2 NaHCO3, 1.25 NaH2PO4 and 11 D-glucose. Coronal brain slices (300 μm) containing the prelimbic cortex (Paxinos and Watson, 5th edition) were cut on a vibratome tissue slicer (Ted Pella Inc., USA) and transferred to incubation chamber, where they were incubated in ACSF solution for at least 1 hr at room temperature before recording. The ACSF solution was constantly bubbled with 95% O2-5% CO2 to maintain pH 7.4.

Whole-cell current-clamp recordings were performed using standard procedures at room temperature. Brain slices were transferred to a submersion-type chamber and perfused constantly (~2 ml/min) with ACSF. Layer 5-6 pyramidal cells were visualized using an Olympus BX51 microscope equipped with IR-DIC optics and an infrared video camera (Qimaging, Canada). Current-clamp recordings were obtained using Axon 200B amplifier, Digidata 1322 A/D converter and pClamp software (Molecular Devices, USA). Recordings were not corrected for the liquid junction potential. Data were discarded if Ra was altered by ~20%. For IPSC recordings, external perfusion solution contained AMPA receptor antagonist CNQX or DNQX (20 μM) and NMDA receptor antagonist APV (50 μM). The pipette solution contained (in mM) 70 K-gluconate, 70 KCl, 20 HEPES, 0.5 CaCl_2_, 5.0 EGTA, 5 Mg-ATP, and its pH was adjusted to 7.2 with KOH. The pipette resistance, as measured in the bath, was typically 2.0-3.0 MΩ. Ion channel blockers used in this study were applied by bath perfusion for at least 10 min unless otherwise noted. To assess the role of HCN channels in GABAergic transmission, the HCN channel blocker ZD7288 was applied after 5~10 min baseline recording. ZD7288-induced changes in GABAergic transmission were measured in the last 3 min of the 15-min perfusion of ZD7288 unless otherwise mentioned.

### Chemicals

All reagents were purchased from the Sigma Chemical Company (St. Louis, MO) with the exceptions of ZD7288 from the Tocris company (UK), ω-Agatoxin IVA, ω-Conotoxin GVIA and pimozide from Alomone Labs (Isreal). All channel blockers were prepared as concentrated stock solutions in distilled water or DMSO and either added immediately to ACSF at working concentrations or stored at −20°C for subsequent utilization.

### Immunostaining

Age-matched male Sprague-Dawley rats were anesthetized with pentobarbital sodium (40 mg/kg, i.p.), and transcardial perfusion was performed with 34°C saline (200 ml) followed by 4% ice-cold paraformaldehyde (PFA) in phosphate-buffered saline (PBS, pH 7.4). Brains were removed and were fixed for 24 h in PFA at 4°C. After that, the brains were put in 30% (w/v) sucrose solution. Coronal sections (35 μm) were cut using a cryostat (Leica CM900, Germany). Brain sections were rinsed with 0.01 M PBS and incubated in a solution of 0.5% Triton-X in PBS for 15 min, followed by normal blocking solution (goat serum, Invitrogen) for 2 h at room temperature. Sequential primary immunolabeling for HCN1, HCN2 or HCN4 was performed using anti-HCN1, HCN2 or HCN4 rabbit antibodies (1:40; Alomone Laboratories, Israel,

Product# APC-056, APC-030, APC-057) [63]. GABAergic neurons were labeled using anti-parvalbumin mouse antibody (PARV-19, 1:1000; Sigma, St. Louis, MO, USA, Product# P3088). All primary antibodies were diluted in goat serum (Invitrogen) and incubated 48 h at 4 °C. After 48-h incubation, the brain sections were rinsed with PBS, and an appropriate secondary antibody was applied. Fluorescent secondary antibodies (whole IgG affinity-purified antibodies: Goat anti-rabbit FITC, Goat anti-mouse Texas Red and Goat anti-mouse FITC; all from Jackson ImmunoResearch Laboratories, West Grove, PA, USA) were applied at a 1:100 dilution in normal blocking serum for 2 h at 4 °C.

### Confocal microscopy

Immunolabeled sections were examined using a confocal laser-scanning microscope system (Leica SP2, Mannheim, Germany). FITC and Texas Red fluorochromes were excited at 488 nm and 543 nm wavelengths, respectively, and the fluorescent emission was collected through BP 505-530 and BP 560-615 filters, respectively. Twelve-bit images were captured at a resolution of 1024×1024 pixels using a 20× objective and at 1024×1024 pixels with a Plan-Apochromat 63/1.4 oil-immersion. Immunoreactivity (IR) was examined under optimal resolution (small pinhole, thin optical slice, and high numerical aperture oil-immersion objective). The pinhole diameter was set to 1.5 airy unit to reduce the influence of cytoplasmic fluorescence as much as possible. Z-sectioning was performed at 0.5-μm intervals, and stacks of optical sections at the *z* axis were acquired. For comparison of the distribution of HCN1, HCN2 and HCN4 channels, each micrograph was captured using the same settings for laser power, pinhole, and photo-multiplier gain. Confocal photomicrographs were processed using Adobe Photoshop (San Jose, CA, USA). No immunolabeling was observed in control slices in which the primary antibody was omitted. The multi-tracking configuration was employed to rule out crosstalk between the fluorescent detection channels.

### Data analysis and statistics

Data are expressed as mean±SEM in all cases. The frequency and amplitude of sIPSCs/mIPSCs were analyzed using the Mini Analysis software package (v8.0, Synaptosoft, Leonia, NJ, http://www.synaptosoft.com). Events above five-fold baseline noise level in amplitude were detected and were used for analysis. Drug-induced changes in cumulative fractions of sIPSC/mIPSC amplitude and inter-event interval were analyzed for statistical significance using the Kolmogorov-Smirnov (K-S) test (Mini Analysis v8.0) and a conservative critical probability level of *p*<0.01. Grouped data were analyzed using paired or unpaired *t*-test for two-group comparison, one-way ANOVA for multi-group comparison, and a critical probability of *p*<0.05 (STATISTICA 6.0, USA).

## Acknowledgements

The present study was supported by grants from the Ministry of Science and Technology of China (2006CB500807; 2011CBA00406 to BML), the National Natural Science Foundation of China (30990263, 30821002, 31121061 to BML, 30700218 to XHZ) and Open Research Fund of the Key Laboratory of Brain Functional Genomics, Ministry of Education (XHZ).

## Competing interests

The authors declare no competing or financial interests.

## Notes

### Competing Interest Statement

The authors have declared no competing interest.

